# The epidemiology and genomics of a virulent emerging fungal pathogen in an Australian reptile

**DOI:** 10.1101/2021.10.22.465237

**Authors:** B. Class, D. Powell, J. Terraube, G. Albery, C. Delmé, S. Bansal, C.H. Frère

**Affiliations:** Global Change Ecology Research Group, University of the Sunshine Coast; Sippy Downs, QLD, Australia; Department of Biology, Georgetown University; Washington DC, United States; School of Biological Sciences, University of Queensland, St Lucia, QLD, Australia

## Abstract

Emerging infectious fungal diseases (EIFDs) represent a major conservation concern worldwide. Here, we provide early insights into the potential threat that *Nannizziopsis barbatae* (*Nb*), a novel EIFD, poses to Australian herpetological biodiversity. First known to the reptile pet trade as a primary pathogen causing untreatable severe dermatomycosis, since 2013, *Nb* has emerged in a growing number of phylogenetically and ecologically distant free-living reptiles across Australia. Observing its emergence in a long-term study population of wild eastern water dragons (*Intellagama lesueurii*), we demonstrate the pathogen’s virulence-related genomic features, within-population spatiotemporal spread, and survival costs, all of which imply that *Nb* could pose a threat to Australian reptiles in the future. Our findings highlight the need to closely monitor this pathogen in Australian ecosystems.

## Introduction

Emerging infectious fungal diseases (EIFDs) pose a serious threat to the conservation of global biodiversity (*1*–*3*) and are responsible for some of the most severe mass mortality events in wild populations (*1*–*4*). Notable examples include chytridiomycosis, which has now impacted over 500 species of amphibians in 54 countries, driving the extinction of 90 species worldwide (*5*); white-nose syndrome, which has resulted in a devastating 75% population decline across bats in Canada and the USA (*6, 7*); and the more recent snake fungal disease (*8*), which poses a significant threat to snake populations in eastern North America (*9*). Whilst EIFDs make up less than three percent of infectious agents reported amongst animal hosts, they are nonetheless responsible for over 70 % of disease-driven population declines and extinctions (*1*).

Members of the fungal genus *Nannizziopsis* are well known to the pet trade as primary pathogens that cause serious cutaneous and systemic fatal disease in a diverse range of reptiles across the world (*10*–*13*). *Nannizziopsis barbatae* was first identified in captivity in 2009 (*14*), and remained confined to captivity until, in 2013, two-free living eastern water dragons (*Intellagama lesueurii*) from locations separated by 30 km across Brisbane (Queensland, Australia) were identified with proliferative dermatitis, necrosis, ulceration and emaciation (*15*). *Nb* has since emerged in a growing number of phylogenetically and ecologically distant free-living lizards (2 x agamid species and 2 x skink species) across Australia (6 sites in Qld, 1 site in NSW and 1 site in WA) (*15*) and is known to cause disease in 9 species (data combined from captive and wild cases, see Table S1). This recent emergence in the wild, followed by a rapid expansion of its geographical distribution and host range, indicate that this fungal pathogen may present a pressing new threat to Australia’s herpetological biodiversity. While we know that *Nb* causes untreatable severe dermatomycosis (*15*), mitigating its impact will require a thorough understanding of its ecology. Taking advantage of its recent emergence in a long-term study population of eastern water dragons (*15*) and using an innovative combination of comparative genomics and spatiotemporal autocorrelation models, we assess the potential threat that *Nb* may pose to Australia’s herpetological biodiversity.

## Results

*Nb* emerged in 2013 in two geographically isolated populations of eastern water dragons in the city of Brisbane (QLD, Australia). One of these populations (Roma Street Parkland, 27°27′46′S, 153°1′11′E) has been monitored with frequent behavioral surveys and yearly catching since 2010. This population comprises 336 individuals on average and behavioral surveys were performed 2-3 times per week along a transect which covered 85% of this population (*16*). During behavioral surveys, individuals’ GPS position were systematically recorded and profile photographs taken to allow later identification based on unique scale patterns (*17*). Disease diagnosis was based on the presence or absence of characteristic skin lesions (*15*), which observers were trained to recognize from season 9 (2018-2019) onwards. Individuals’ disease status for earlier years was hence determined retrospectively using photos from catching and, when not available, behavioral surveys. Once diagnosed, individuals were assumed to remain diseased even when not caught again. Amongst the diseased individuals repeatedly captured between February 2020 and August 2021, some individuals (20/61) showed a reduction in the severity of their lesions, although this reduction was mainly observed in individuals exhibiting mild lesions (15/20, Data S1). Using this decade-long individual-based data, we found that the disease prevalence has continuously increased throughout the population since *Nb*’s emergence. Starting with one individual in 2013, a total of 158 individuals have now presented with clinical signs of the disease (n =1221 for field seasons 3-11) and in the last field season (2020-2021), the prevalence was 26.4% (95%CI: 24.2-28.4, Fig. S1, Table S2). The majority (96.7%) of these individuals were adults, and males (58.3%). Only five juveniles were found with clinical signs of the disease (0.07 to 5.5% of juveniles) between late 2018 to early 2021, despite juveniles representing on average 17.8% of observed individuals during these years.

### N. barbatae shares genomic characteristics with other fungal pathogens

*Nb* is a member of the Onygenales, an order of fungi that are able to degrade keratin, the main component of the vertebrate outer skin layer. Some members of this order are important primary pathogens of animals and humans and recent comparative genomic studies have helped resolve differences in gene content between pathogenic and non-pathogenic species (*18*). With this in mind, we performed a comparative whole-genome analysis incorporating the full set of genes of *Nb* (8,012 predicted protein-coding sequences) together with 16 other species of fungi (Table 1) to uncover genomic features likely to contribute to *Nb*’s pathogenicity.

**Table 1.**
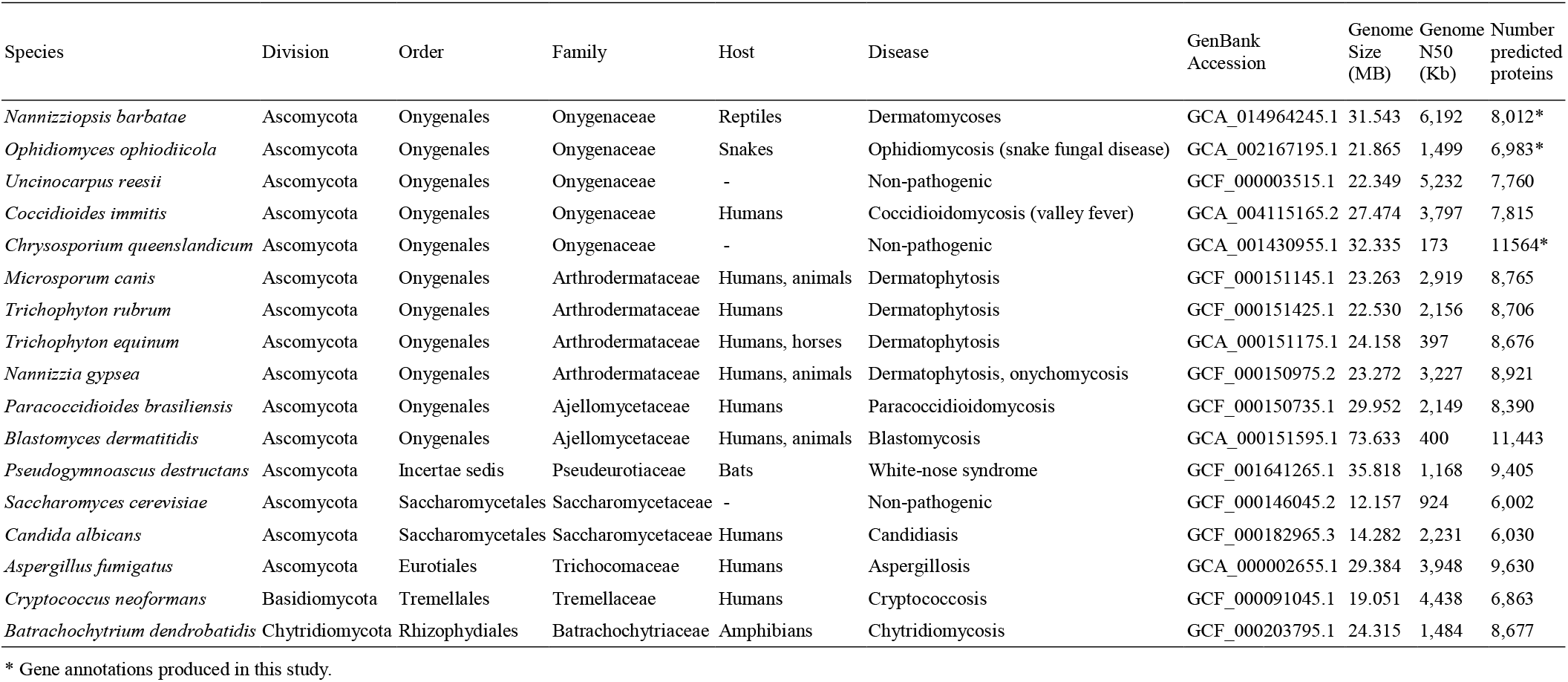
Details of the fungal species used in the comparative analysis.

First, we identified that the *Nb* genome contains a gene repertoire rich in proteases, known to increase fungal virulence (*19*), and shares similarities with other pathogenic Onygenaceae (Fig. 1A). Most notable is an expansion of trypsin domain-containing genes (PF00089) (Fig. 1B) found only in *Nb* (7 genes) and the fungus causing snake fungal disease, *O. ophidiicola* (29 genes). Both of these species are capable of primary infection in reptiles (*9, 20*) suggesting a role for this gene family in influencing host range. *Nb* has a degradome that bears resemblance to important dermatophytes and the enrichment of proteases including, subtilase (PF00082) and deuterolysin (PF02102), suggest extensive proteolytic capacity. Second, *Nb* has a large number of protein kinase domain-containing genes (PF00069) which may contribute to its capacity to infect a broad range of reptile taxa (*15*). Last, we identified a higher number of LysM domain-containing genes (PF01476) in the *Nb* genome than most of the other fungi in this analysis. Together, these characteristics of the *Nb* genome highlight factors which may be key to determining its propensity to infect herpetofauna.

**Fig. 1.**
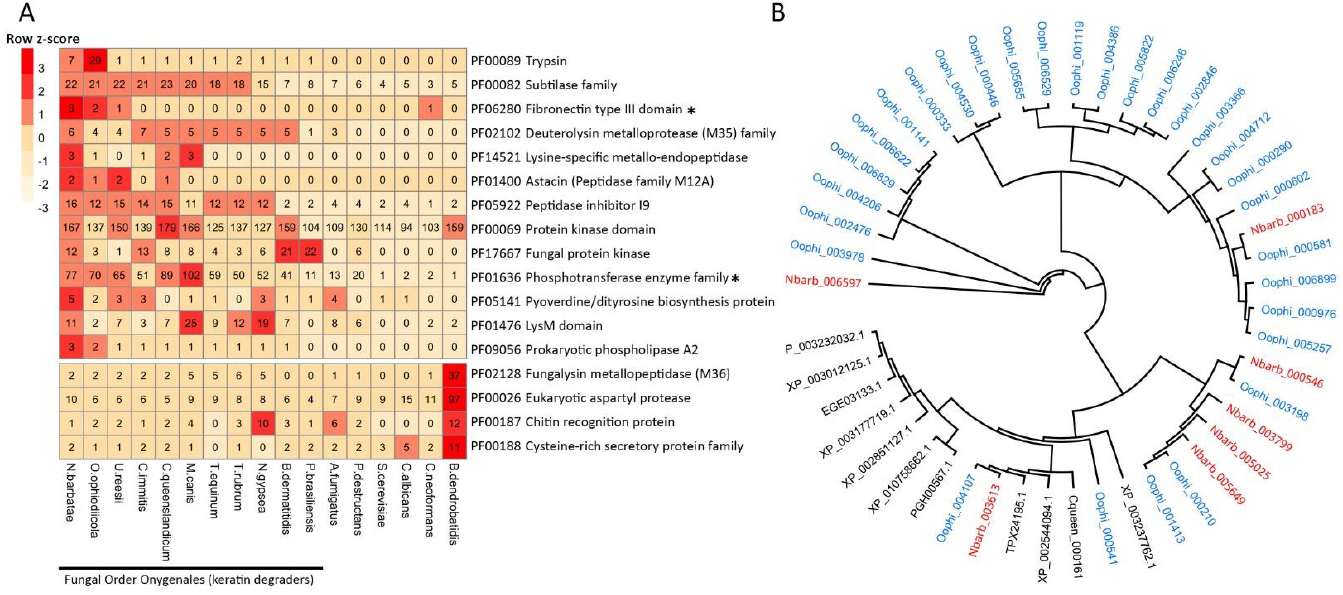
Comparative genomic analysis of *N. barbatae* with other fungal species. (**A**) Gene family size comparison of putative proteases and other proteins implicated in fungal virulence. Protein families significantly expanded in *N. barbatae* are marked with *. (**B**) Phylogenetic relationship of the trypsin domain-containing protein sequences identified in this study, red text, *N. barbatae* proteins; blue text, *O. ophidiicola* proteins.

### N. barbatae infection is spatially structured within the population

To investigate the phenotypic and spatiotemporal predictors of fungal infection, we constructed spatiotemporal autocorrelation models using the Integrated Nested Laplace Algorithm. Comparison of disease prevalence models (Fig. S2) provided strong support for spatial structuring (ΔDIC=-117.24 relative to the base model), but relatively little evidence for spatiotemporal structuring (ΔDIC=-4.11 relative to the spatial model) (Fig. S2). That is, disease prevalence varied more spatially (assuming no time effect) than spatiotemporally. Indeed, spatial effects were strongly correlated across field seasons (rho>0.9) and disease prevalence has remained consistently higher in the East (up to 33%) compared to the West (<10%, Fig. 2). Models also showed lower prevalence in juveniles than adults, and in females compared to males (Fig. S2).

**Fig. 2.**
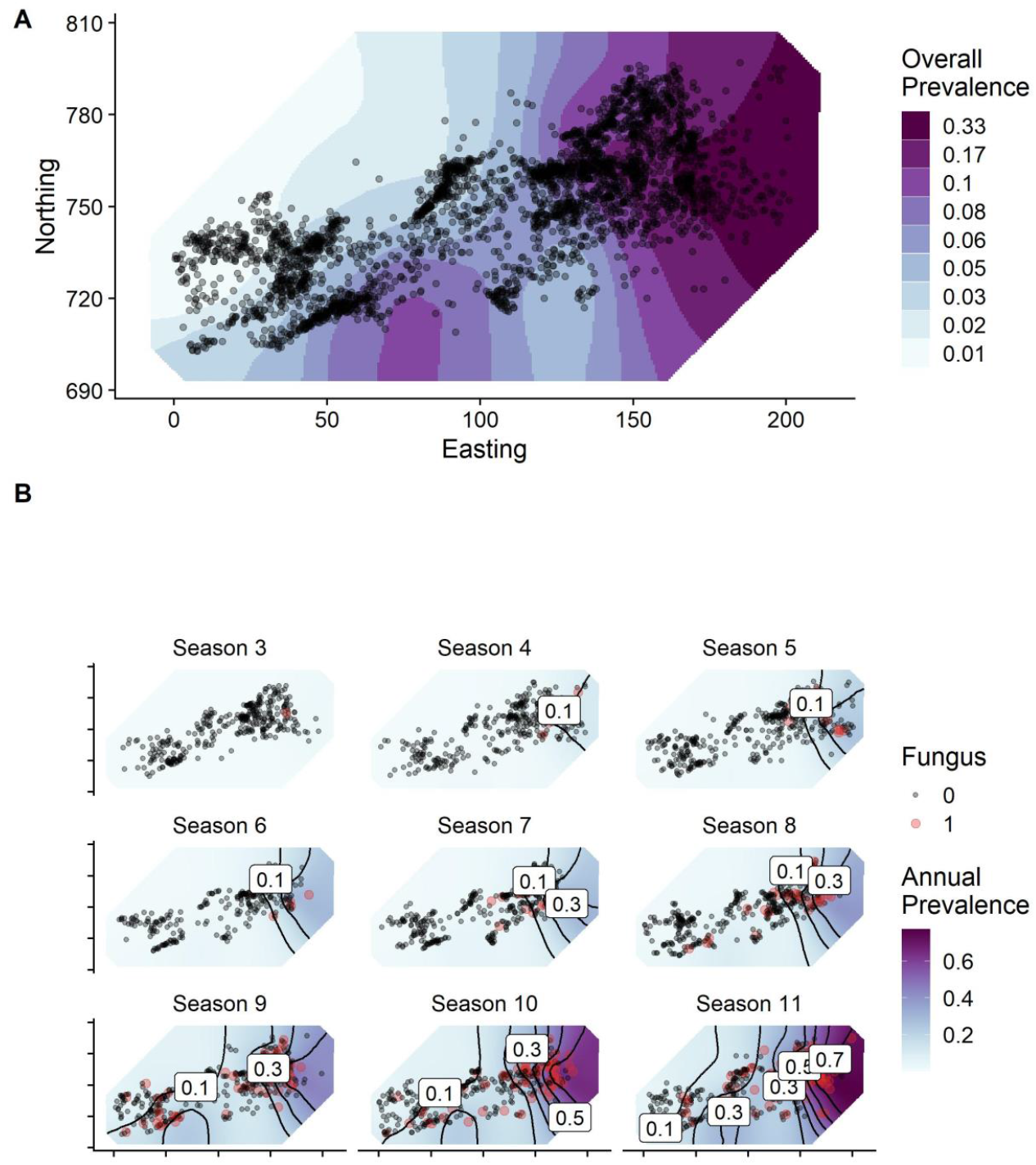
Spatial distribution and temporal spread of *N. barbatae* infection within the population. These are represented as the spatial distribution of the spatial random effect from the Spatial model (**A**) and the annually stratified Spatiotemporal model (**B**), respectively. The spatial effects were estimated using a stochastic partial differentiation equation (SPDE) in an integrated nested Laplace approximation (INLA) model. Adding these SPDE components substantially improved model fit. In (A), points represent individuals’ average annual locations. The axes in (B) are identical to those in (A), with the labels removed for plotting clarity. In (A), the spatial effect is categorized into eight quantiles to facilitate visualization over a range of prevalence values.

### N. barbatae infection is associated with survival costs

To investigate the survival costs of infection, we fitted a binomial survival model, where survival of an individual was coded based on whether they were observed in any subsequent year. The full population model showed effects of cohort, field season and sex on the yearly probability of survival but failed to detect any effect of the disease (Fig. 3A). In contrast, randomly subsampling diseased and non-diseased individuals from matching cohorts and accounting for age (number of days in the population), sex, and field season in subsequent models revealed a small but significant individual survival costs of the disease (Fig. 3B-D). All subsampled models found a significant effect of the disease (Fig. 3B); the overall mean survival cost was 12% (Fig. 3C), so that the mean predicted annual survival of diseased individuals was 74% compared to 86% for non-diseased individuals (Fig. 3D). Controlling for spatial autocorrelation did not improve model fit (ΔDIC>-2 relative to the base model), demonstrating that this survival cost did not vary spatially.

**Fig. 3.**
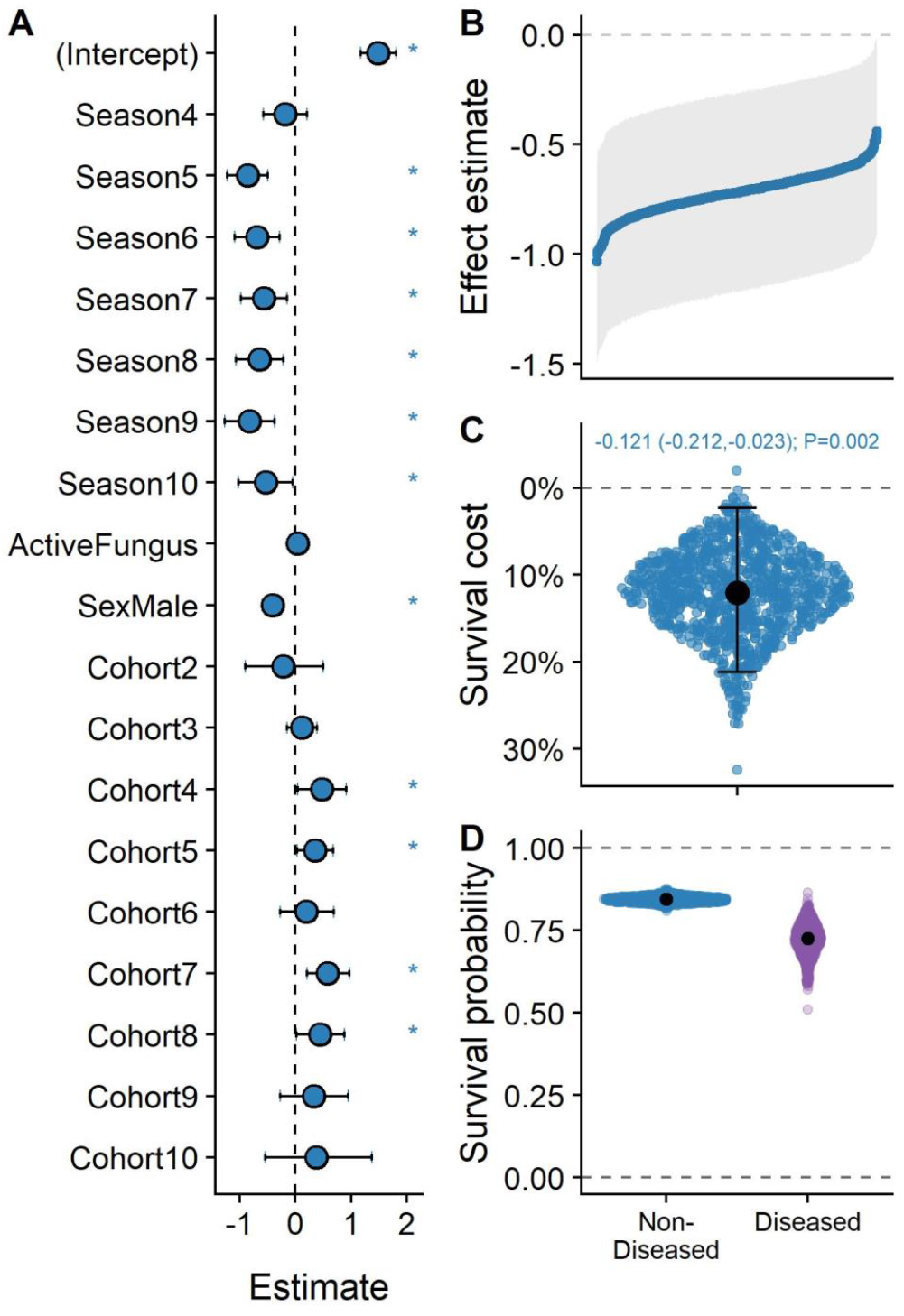
Survival effects of *N. barbatae* infection in the population. Survival effects of the disease are estimated using a full-population model (**A**), and an approach combining model uncertainty and subsampling regime uncertainty (**B, C, D**). The second approach provided estimates of survival effects across all subsamples (B) and survival costs (C) and probabilities (D) of diseased vs. non-diseased individuals. The large black points represent means across all 1000 replicates. The text at the top of panel C displays the effect estimate for the survival cost across all models, with 95% credibility intervals in brackets and the P value. The error bars represent the 95% credibility intervals.

## Discussion

Gaining early insights into disease virulence, spatiotemporal spread, and survival costs is particularly urgent in the case of novel emerging infections that have the potential to severely threaten biodiversity. Yet such data are challenging to obtain in the wild, greatly impeding our abilities to predict and mitigate the impact of infections on wildlife. Our study investigates within-population spread and impacts of *Nannizziopsis barbatae*, a novel emerging infectious fungal disease which should give us cause for vigilance.

### Genomic signatures of pathogenicity

We show that the free-living *Nb* genome sampled from our long-term study population of eastern water dragon contains many gene families implicated in fungal pathogenicity, including several proteases, protein kinases, and LysM effector proteins. Virulence in wildlife fungal pathogens has often been associated with expansions of protease gene repertoires and their expression (e.g. chytrid (*21*); WNS (*22*–*24*)). *Nb* has a gene repertoire rich in proteases with features similar to those identified in other wildlife fungal pathogens such as snake fungal disease (e.g. trypsin domain-containing genes) and chytrid fungus (e.g. M36 metalloproteases). Additionally, a comprehensive and novel repertoire of protein kinases can provide fungal species with plasticity in occupying different ecological niches and responding to environmental change (*25, 26*). LysM effector proteins may contribute to fungal virulence by suppressing the host immune system response via interactions with chitin (*27*). Comparative studies strongly suggest an association with the enrichment of LysM domain-containing genes and virulence in the keratin-degrading dermatophytes (*26*). Furthermore, chitin-binding CBM18 gene family proteins (PF00187) are also found expanded in *B. dendrobatidis*, and are thought to play a role in evasion of the amphibian host immune response (*26*). While understanding the molecular mechanisms of this pathogen is central to mitigating its impact on wildlife, the exact source of Nb, its current free-living genetic diversity, and its mode of introduction into our dragon population remain unknown. Genomic resources for Nannizziopsis spp. will enable the development of tools to answer these questions.

The emergence of this disease urgently necessitates the identification of its origin to better understand and thus predict the impact it will likely have on the Australian herpetofauna. For instance, it is critical that we determine whether or not we are dealing with a novel pathogen and thus naïve hosts, or whether the population has had historical exposure to the pathogen.

### Within-population spread

Even though central to the forecasting and control of wildlife disease management, quantifying the contribution of different transmission pathways of a pathogen is notoriously challenging to achieve in nature (*28*). Using an intensively-studied lizard population, we provide a much-needed early assessment of *Nb*’s spatiotemporal spread since its emergence in 2013. We show that the disease has spread relatively rapidly across an increasing portion of the population, providing the first likely evidence for within-population *Nb* transmission in the wild. From a single individual dragon identified with *Nb*-like clinical signs in 2013, more than 150 individuals have displayed apparent clinical signs of *Nb* and the disease prevalence has reached 26.4% of the population. Worryingly, prevalence of the disease has been continuously increasing since 2016 and shows no signs of slowing down. Although it is unclear what the transmission route of this pathogen is in this population (e.g. physical contact or environmental latency), our results show that over the years, the disease prevalence has remained higher in the eastern part of the park than in the western area, which could be due to spatial variation in environmental factors influencing the pathogen’s survival, transmission, or virulence (*29*). Analyses of the dragons’ spatial and social behaviors coupled with molecular diagnostics capabilities will help identify transmission routes, predict geographic spread of the pathogen, and inform potential future interventions (*30*).

### Low but detectable survival costs

Survival costs of *Nb* infection were detectable at an individual level. Although individuals showing clinical signs of the disease varied in their survival costs, they were still relatively likely to survive from one year to the next (>70% chance), demonstrating that adults are relatively tolerant to the pathogen and can carry it for multiple years once skin lesions become apparent. Although the disease has been shown to be incurable in captive reptiles (*15*), some rare individuals in this population showed reductions in the severity of their lesions (Data S1) and we are yet to determine whether individual diseased dragons can entirely clear the infection (as shown in chytridiomycosis (*31*) and white-nose syndrome (*32*)). Additionally, we were only able to detect survival costs when we subsampled our dataset to cohort-matched (age and sex) diseased and non-diseased individuals, thereby reducing extraneous variation in survival probability. Evidence for individual survival costs remains similarly equivocal for other EIFDs, some of which have been studied for much longer than *Nb* (*31*–*33*). We also acknowledge some uncertainty in our estimates of survival costs due to potential errors in diagnosis, our visual assessment being particularly prone to miss asymptomatic or cryptic infections in the population. Additionally, because lesions are easier to observe in caught individuals, this underestimation may be particularly severe for individuals or classes of individuals that were less likely to be caught (e.g. juveniles). Taken together, these facts imply a general difficulty detecting survival costs of fungal pathogens in long-lived reptiles.

Despite identifying individual-level costs of infection, predicting *Nb*’s impacts on population dynamics remains difficult. Such uncertainty is likewise common to other EIFDs, as some populations affected by chytridiomycosis and white nose syndrome have not declined systematically (*44, 45*). Predicting *Nb*’s impacts on the viability of our studied population of eastern water dragons will require key information about: i) *Nb*’s prevalence and survival costs at different life stages (*36, 37*), which should be achieved with higher certainty through the use of molecular diagnosis; ii) *Nb*’s potential reproduction costs, as was documented for snake fungal disease (*38*) and chytridiomycosis (*39*); iii) the mechanisms underlying *Nb*’s spatiotemporal spread. In addition, assessing whether the pathogen’s transmission dynamics are density-dependent might be crucial to understanding whether the epidemic will become self-limiting (*40*).

### A novel threat for the Australian biodiversity?

EIFDs constitute an increasing cause for concern regarding global health, food security and biodiversity conservation (*1*). With *Nb*, we may be witnessing the early days of a novel fungal threat to Australian herpetological biodiversity. While other infamous EIFDs with global impacts on wild animal populations were only reported after mass mortality events had already occurred (*41, 42*), we have the unique opportunity to monitor the emergence of this pathogen and take action early enough to limit its spread. Although the origin and long-term population impacts of *Nb* remain unknown, its genomic similarity with other pathogenic EIFDs, capacity to spread in the wild, and detectable survivals costs, combined with its repeated emergence across the country and broad host range, highlight the critical need to closely monitor this pathogen in Australian ecosystems.

## Materials and Methods

### Study system

The population of Eastern water dragons has been monitored since 2010, with frequent behavioral surveys and regular catching during the active season, (i.e. early September to late April). During behavioral surveys (2-3 times per week), individuals’ behavior and GPS position were recorded and photographs taken to allow later identification based on unique scale patterns (see *(17)*). Individuals were also caught during 1-2 weeks catching events in the years 2013, 2014, and yearly since 2016. Morphometric measurements, head and body photographs and DNA samples (blood or tip of the tail) were taken, and unique PIT-tags were inserted in their right upper hind leg. EWD are sexually dimorphic, males being overall larger than females, with more developed jaw and dorsal crest and a red ventral coloration (*43*). Age class (adult vs. juvenile) was determined for each breeding season using a combination of approaches (snout-vent length when individuals were caught; general appearance when individuals were not caught) and taking into account individuals’ observation history (individuals being considered adults after 3 years (*43*)). Disease diagnosis was based on the presence or absence of characteristic skin lesions (*15*), which observers were trained to recognize from season 9 (2018-2019) onwards. Individuals’ disease status before season 9 was hence determined retrospectively using pictures from behavioral surveys and catching (75-100% for the latter). From season 9 onwards, disease status was assessed directly in the field during behavioral surveys and catching (65-92% for the latter). From February 2020 onwards, disease severity was rated for captured individuals using scores ranging from 0 (no lesions, not diseased) to 5 (severely diseased, Table S3).

### Genome annotation and comparative analysis

The genomes of *N. barbatae, O. ophidiicola* and *C. queenslandicum* were annotated using the Funannotate (v1.7.4) gene prediction pipeline (https://funannotate.readthedocs.io/). Genomes were firstly screened for repeats using custom generated databases for each species using RepeatModeler (v2.0.1) and masked using RepeatMasker (v4.1.0; http://www.repeatmasker.org). Repeat masked assemblies were then cleaned and sorted before initial gene prediction using GeneMark-ES (v4.65) (*44*). Protein sequences from high-quality fungal genomes used in this study were used for protein-to-genome alignments as evidence for gene predictors AUGUSTUS (4*5*), SNAP(4*6*), and Glimmer (*47*) before being passed to EVM (4*8*) to build consensus gene structures. All other predicted protein sequences were downloaded directly from GenBank (Table 1). The newly annotated gene models were evaluated for completeness using BUSCO (v5) (4*9*) in protein mode against the ascomycota_odb10 database (Table S4).

Gene families within each fungal genome were identified from searches of the protein-coding sequences for Pfam (*50*) domains to assign gene function. We used HMMER (v3.1) (*51*) (hmmscan) to search the Pfam A database (release 32.0) for 4312 different domains of 16 different species of fungi. To test for significantly expanded gene families, a Fisher’s exact test was then conducted iteratively using R (*52*), comparing the number of counts in Pfam families found in an individual genome, normalised by the total gene count for that species, against the background, which we defined as the average of the counts in the remaining species. Multiple testing corrections were done using the FDR method in R for all calculated *p*-values. A Pfam domain was considered expanded if it showed a corrected *p*-value < 0.05. Counts of each domain were collated for each species with domains that occurred multiple times in a protein sequence being counted only once. Heatmap was generated using the package pheatmap with data normalised using the scale function in R. Protein sequences were aligned using Muscle (v3.8.425) (*53*) and phylogenetic inference made using FastTree (v2.1.12) (*54*) built in to the commercially available Geneious Prime (v2021.1.1) software.

### Drivers and spatiotemporal dynamics of infection

To investigate the phenotypic and spatiotemporal predictors of fungal infection, we constructed spatiotemporal autocorrelation models using the Integrated Nested Laplace Algorithm, implemented in the ‘inla’ package in R (*55*). These models fitted binary fungal infection as a response variable, where an individual was coded as a 1 if it had previously been diagnosed with fungal infection, and a 0 otherwise. All covariates were categorical, and included Age class (3 levels: Adult, Juvenile, and Unknown); Sex (2 levels: Female and Male); Field season (9 levels: one for each sampling year 2012-2021). The model used a binomial logit error distribution:

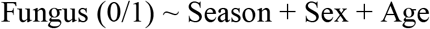

We first fitted these fixed effects as a “Base” model. To investigate spatiotemporal patterns of infection, we then added Stochastic Partial Differentiation Equation (SPDE) random effects using individuals’ mean map locations in a given season (“annual centroids”). This random effect models two-dimensional patterns of the response variable based on distances between individuals using Matérn correlation (*56, 57*) The “Spatial” model used a static field, where the spatial distribution of infection was modelled to be unchanging across the study period; the “Spatiotemporal” model allowed this field to change from year to year, using an autoregression (AR1) correlation across years, to examine how the infection’s distribution changed over the course of the study period. We compared these three models using deviance information criterion (DIC) as a measure of model fit to investigate whether spatiotemporal correlation significantly improved the model.

### Survival costs

To investigate the survival costs of infection, we fitted a binomial survival model, where survival was coded based on whether the individual was seen in a subsequent year (we hence excluded the most recent year, 2019). The model was specified as follows:

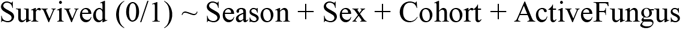

Following this model, we used a subsampling routine that allowed us to reduce extraneous variation in survival by compensating for the low proportion of infected individuals in the study period (120/1151=10.4%) and for the unknown age of infected individuals. We 1) assigned each individual a cohort based on the first season that they were observed in the population; 2) selected the 101 individuals that were ever observed with an infection between 2012 and 2018; and 3) age matched each diseased individual with a random non-diseased individual from their cohort. Between 2018 and 2020, six diseased individuals that were caught in a very poor condition were euthanized. These individuals were hence excluded from these analyses. Having subsampled the population, we then ran the same model as before. This protocol was repeated 1000 times to ensure an even and different selection of non-diseased individuals and survival effect estimates.

We summarized the findings from these models by predicting survival probability for each individual and comparing these values between uninfected and infected individuals. To produce conservative estimates, we randomly drew one effect estimate from each model’s fungal effect estimate posterior distribution and used these estimates to predict the survival probability for all infected and uninfected individuals. We then took the mean survival probability for these groups of individuals and subtracted the infected individuals’ survival probability from those of the uninfected individuals to estimate a survival cost of infection.

## Acknowledgments

We acknowledge the Turrbal and Yugara people, as the First Nations owners of the lands where our study site sits. We pay respect to their Elders, lores, customs and creation spirits. In addition, we would like to thank the students and volunteers that have contributed to the data collection as well as the staff and management of Roma Street Parkland for their ongoing support.

## Author contributions

Conceptualization: BC, JT, CF, DP, GA, SB

Data curation: BC, CD, CF

Methodology: GA, DP, BC

Investigation: BC, DP, JT, CF, CD

Visualization: DP, GA

Funding acquisition: CF

Project administration: CF

Supervision: CF, SB

Writing – original draft: BC, JT, CF

Writing – review & editing: BC, JT, CF, DP, GA, SB, CD

## Data and materials availability

Genome annotation data, individual data and R code used for statistical analyses are available from Figshare using the following link https://doi.org/10.6084/m9.figshare.16599245.

## Supplementary Materials

Figs. S1 to S2

Table S1 to S4

Data S1

