## Supplemental figures and tables for "The epidemiology and genomics of a virulent emerging fungal pathogen in an Australian reptile"

### Supplementary Materials for: The epidemiology and genomics of a virulent emerging fungal pathogen in an Australian reptile

B. Class, D. Powell, J. Terraube, G. Albery, C. Delmé, S. Bansal, C.H. Frère

**This file includes:**

Figs. S1 to S2
Tables S1 to S4
Captions for Data S1

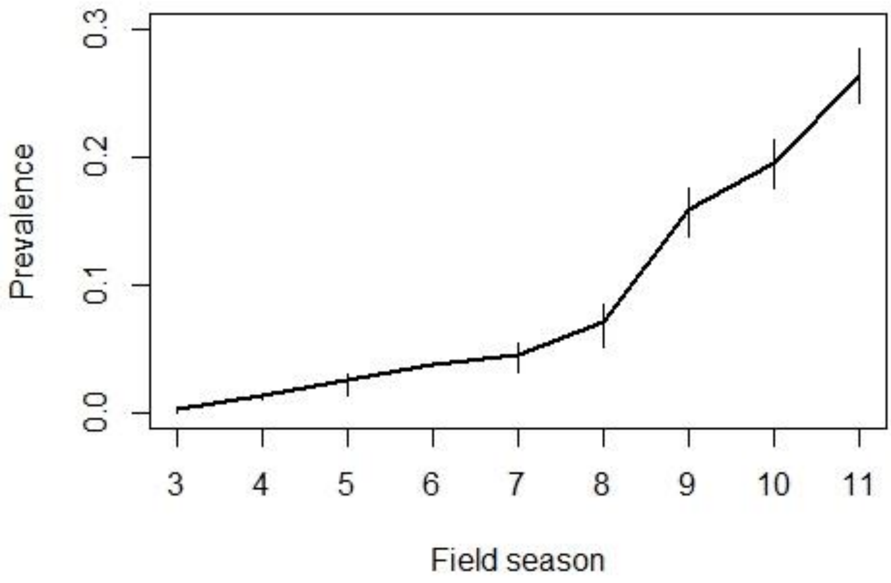


**Fig. S1.** Prevalence (proportion of diseased individuals) observed for each field season from 2012-2013 (season 3) to 2020-2021 (season 11). Error bars represent sampling uncertainty and were obtained by randomly sampling 295 individuals (minimal number of unique individuals in season 6) in each field season 100 times.

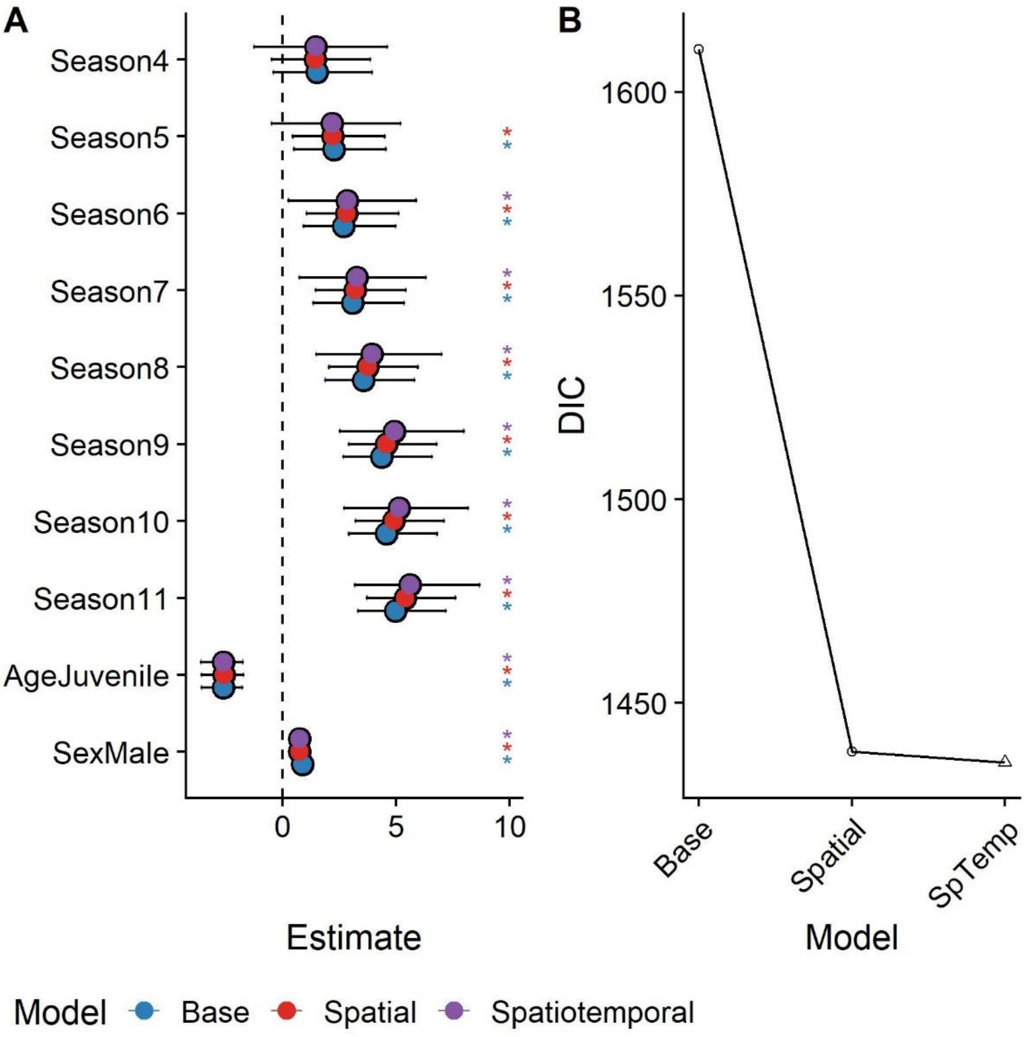


**Fig. S2.** Model output (A) and DIC change (B) for the base, spatial, and spatiotemporal models of infection prevalence.

**Table S1.** Species found to be infected by *N. barbatae* in the wild and in captivity.

| **Wild *N. barbatae* cases** | **Captive *N. barbatae* cases** |
|---|---|
| Eastern water dragons (*Intelllagama lesueurii*) | Centralian blue tongue skink (*Tiliqua multifasciata)* |
| Eastern blue tongue skink (*Tiliqua scincoides scincoides*) | Pygmy spiny-tailed skink (*Egernia depressa)* |
| Tommy roundhead dragon (*Diporiphora australis)* | She-oak skink (*Cyclodomorphus casuarinae)* |
| Singleback skink (*Tiliqua rugosa)* | Central bearded dragons (*Pogona vitticeps*) |
| | Monitor lizard (*Varanidae*) |

**Table S2.** Observed *Nb* prevalence for each field season and its bootstrapped confidence interval. Abbreviations: n.dis is the number of diseased individuals, n the total number of individuals encountered for each field season, prev is the observed prevalence (n.dis/n) and 2.5%,50% 97.5%, are the bootstrapped prevalence's 95% confidence interval and median.

| Field season | n.dis | n | prev | 2.5% | 50% | 97.5% |
|---|---|---|---|---|---|---|
| 3 | 1 | 344 | 0.003 | 0.000 | 0.003 | 0.003 |
| 4 | 4 | 305 | 0.013 | 0.010 | 0.014 | 0.014 |
| 5 | 10 | 400 | 0.025 | 0.014 | 0.024 | 0.031 |
| 6 | 11 | 295 | 0.037 | 0.037 | 0.037 | 0.037 |
| 7 | 17 | 381 | 0.045 | 0.032 | 0.046 | 0.054 |
| 8 | 32 | 449 | 0.071 | 0.051 | 0.071 | 0.085 |
| 9 | 63 | 396 | 0.159 | 0.137 | 0.161 | 0.176 |
| 10 | 71 | 364 | 0.195 | 0.176 | 0.195 | 0.214 |
| 11 | 99 | 375 | 0.264 | 0.242 | 0.264 | 0.285 |

**Table S3.** Scale used to rate disease severity in Eastern water dragons when caught in the field.

| **Individual Rating** | **Clinical Signs – Fungal Disease** |
|---|---|
| 0 – No obvious lesions | Normal, no skin lesions observed (e.g., only injury observed OR unsure of skin condition) |
| 1 – Mild | One to three focal skin lesions ≤ 5mm diameter |
| 2 – Mild/Moderate | Four to five focal skin lesions ≤ 5mm diameter, or one lesion 5-10mm in diameter |
| 3 – Moderate | Up to three lesions 5-10mm diameter, or one to two skin lesions 10-20mm in diameter |
| 4 – Moderate/Severe | Four+ skin lesions 10-20mm in diameter with roughly 5-10% of the skin surface affected |
| 5 – Severe | More than 10% of total skin surface affected, or 5-10% affected and in poor condition |

**Table S4**. Protein-level completeness estimates using the BUSCO tool for the genomes of the three fungal species annotated in this study.

| | ***N. barbatae*** | ***O. ophidiicola*** | ***C. queenslandicum*** |
|---|---|---|---|
| Complete BUSCOs | 1667 | 1663 | 1646 |
| Complete and single-copy BUSCOs | 1666 | 1661 | 1645 |
| Complete and duplicated BUSCOs | 1 | 2 | 1 |
| Fragmented BUSCOs | 6 | 8 | 14 |
| Missing BUSCOs | 33 | 35 | 46 |
| Total BUSCO groups searched | 1706 | 1706 | 1706 |

**Data S1. (separate file)** Identity, date, and *Nb* infection severity of individuals repeatedly caught between February 2020 (start of systematic disease severity recording) and August 2021. Severity was visually assessed during catching, using a pre-defined scale ranging from 0 (no lesions) to 5 (severely diseased).
